# Regulation of nucleolar dominance in *Drosophila melanogaster*

**DOI:** 10.1101/690198

**Authors:** Natalie Warsinger-Pepe, Duojia Li, Yukiko M. Yamashita

**Affiliations:** Department of Molecular and Integrative Physiology, University of Michigan, Ann Arbor; Life Sciences Institute, University of Michigan, Ann Arbor; Molecular, Cellular and Developmental Biology, University of Michigan, Ann Arbor; Cell and Developmental Biology, University of Michigan, Ann Arbor; Howard Hughes Medical Institute, University of Michigan, Ann Arbor

**Author notes:** Corresponding author information: Yukiko Yamashita, Life Sciences Institute, Room 5183, 210 Washtenaw Avenue, Ann Arbor, MI, 48109-2216.

**Keywords:** Nucleolar dominance, rDNA, Drosophila

## Abstract

In eukaryotic genomes, ribosomal RNA (rRNA) genes exist as tandemly repeated clusters, forming ribosomal DNA (rDNA) loci. Each rDNA locus typically contains hundreds of rRNA genes to meet the high demand of ribosome biogenesis. Nucleolar dominance is a phenomenon, whereby individual rDNA loci are entirely silenced or transcribed, and is believed to be a mechanism to control rRNA dosage. Nucleolar dominance was originally noted to occur in interspecies hybrids, and has been shown to occur within a species (i.e. non-hybrid contexts). However, studying nucleolar dominance within a species has been challenging due to the highly homogenous sequence across rDNA loci. By utilizing single nucleotide polymorphisms (SNPs) between X rDNA vs. Y rDNA loci in males, as well as sequence variations between two X rDNA loci in females, we conducted a thorough characterization of nucleolar dominance throughout development of *D. melanogaster*. We demonstrate that nucleolar dominance is a developmentally-regulated program, where Y rDNA dominance is established during male embryogenesis, whereas females normally do not exhibit dominance between two X rDNA loci. By utilizing various chromosomal complements (e.g. X/Y, X/X, X/X/Y) and a chromosome rearrangement, we show that Y chromosome rDNA likely contains *cis* elements that dictate its dominance over the X chromosome rDNA. Our study begins to reveal the mechanisms underlying the selection of rDNA loci for activation/silencing in nucleolar dominance.

## Introduction

Ribosomal DNA (rDNA), genes encoding the catalytic RNA components of ribosomes, is highly repetitive (100s to 1000s of copies) and often exists as multiple loci on separate chromosomes (e.g. 2 loci in *Drosophila melanogaster*, 4 in *Arabidopsis*, 6 in *Mus musculus*, 10 in *Homo sapiens* per diploid genome) (Long and Dawid 1980, Pontes et al. 2004). This expansive copy number may come to no surprise, considering that the transcription of rDNA accounts for ∼60% of the total transcription of a metabolically active cell (Moss and Stefanovsky 2002). The regulation of ribosomal RNA (rRNA) expression is critically important for adjusting cellular energetic expenditure: when nutrients are low rRNA synthesis is downregulated, whereas the opposite occurs when nutrients are high or growth rate is increased (e.g. in cancer) (Smetana and Busch 1964, Busch et al. 1979, Ghoshal et al. 2004, Grewal et al. 2005, Murayama et al. 2008, Aldrich and Maggert 2015). Accordingly, transcription of rRNA is expected to require precise regulation.

A phenomenon called nucleolar dominance, whereby individual rDNA loci are either entirely expressed or silenced, is proposed to be a mechanism that regulates the dosage of rRNA (Preuss and Pikaard 2007). Nucleolar dominance has been noted to be one of the largest epigenetic mechanisms, second only to X-inactivation in eutherian mammals (Pikaard 2000). Nucleolar dominance was originally discovered in interspecies hybrids (i.e. *Xenopus* hybrids (Cassidy and Blackler 1974), *Arabidopsis* hybrids (Chen et al. 1998), *Drosophila* hybrids (Durica and Krider 1977), and mouse-human hybrid cell lines (Croce et al. 1977)), where rDNA loci inherited from one species are preferentially expressed and those from the other are silenced. Later, nucleolar dominance was shown to occur within a species (i.e. non-hybrid context) (Lawrence et al. 2004, Greil and Ahmad 2012, Zhou et al. 2012), indicating that nucleolar dominance is a mechanism to regulate rRNA expression/dosage instead of a result of interspecies incompatibility.

Nucleolar dominance has been thoroughly studied in *Arabidopsis*, both in *A. suecica* (the interspecies hybrid between *A. thaliana* crossed to *A. arenosa)* as well as non-hybrid *A. thaliana* (Pontes et al. 2007, Earley et al. 2010). In both cases, nucleolar dominance is gradually established during development, where seedling cotyledons express rRNA from all rDNA loci (i.e. ‘co-dominance’), transitioning to preferential expression of certain loci in mature tissues (Pontes et al. 2007, Earley et al. 2010). Several mechanisms have been shown to mediate the silencing of chosen rDNA loci, including small interfering RNAs (siRNAs) (Pontes et al. 2006, Preuss et al. 2008), DNA methylation (Chen et al. 1998, Lawrence et al. 2004, Pontes et al. 2006, Preuss et al. 2008, Costa-Nunes et al. 2010, Earley et al. 2010), histone methylation (Earley et al. 2010, Pontvianne et al. 2012), and histone deacetylation (Probst et al. 2004, Earley et al. 2006, Earley et al. 2010). These mechanisms reveal how the large-scale silencing of rDNA is implemented to achieve nucleolar dominance, however, what factor(s) influence the choice of which rDNA loci are silenced or activated remain elusive.

Nucleolar dominance is likely a wide-spread phenomenon across many species. For example, only a subset of rDNA loci are transcribed in human cell lines (Roussel et al. 1996) and human lymphocytes (Roussel et al. 1996, Heliot et al. 2000), implying that these cells also undergo nucleolar dominance. Nucleolar dominance was found to occur in *D. melanogaster* larval neuroblasts, where rDNA on the Y chromosome (‘Y rDNA’) dominates over rDNA on the X chromosome (‘X rDNA’), based on transcription-dependent deposition of GFP-tagged histone H3.3 onto the active rDNA locus (i.e. the Y rDNA locus) (Greil and Ahmad 2012). This method relied on readily available mitotic chromosome spreads, leaving the assessment of nucleolar dominance in other cell types elusive. Recently, we adapted a single nucleotide polymorphism (SNP) RNA fluorescent *in situ* hybridization (SNP *in situ)* protocol and showed that nucleolar dominance (Y rDNA dominance) also occurs in male germline stem cells (Levesque et al. 2013, Lu et al. 2018). This method utilizes SNPs between X rDNA vs. Y rDNA to differentially label their products (X-vs. Y-derived rRNA), allowing assessment of nucleolar dominance without requiring mitotic chromosome spreads.

In this study, we utilized SNP *in situ* to comprehensively examine the state of nucleolar dominance in *D. melanogaster* during development and across different tissues. We show that nucleolar dominance in *D. melanogaster* is gradually established during development, similar to the observations in *A. thaliana*, supporting the notion that nucleolar dominance is a regulatory mechanism that occurs in non-hybrid organisms. We have further examined the state of nucleolar dominance between two X rDNA loci in females by isolating X rDNA with distinct sequences that enables RNA *in situ* hybridization to distinguish transcripts from two X rDNA loci. Our results show that the two X rDNA loci in females exhibit co-dominance in essentially all tissues, expanding the previous finding of co-dominance in female larval neuroblasts (Greil and Ahmad 2012). Moreover, by utilizing the various karyotypes (e.g. X/X females, X/Y males, vs. X/X/Y females) and a chromosome rearrangement strain, we show that Y chromosome element(s) (within Y rDNA as well as non-rDNA element(s) of the Y chromosome) may aid in the ‘choice’ mechanism that preferentially activates the Y rDNA locus and/or silences the X rDNA locus. These results provide insights into how specific rDNA loci may be preferentially transcribed or silenced, and will provide the foundation for future studies aimed at understanding how rDNA loci are chosen for activation/silencing to achieve nucleolar dominance.

## Materials and Methods

### Fly husbandry and strains

All fly stocks (See Reagent Table) were raised on standard Bloomington medium at room temperature. Unless otherwise stated, all flies used for wild-type experiments were the standard lab wild-type strain *y^1^w^1^*, referred to as yw, that contains the X and Y chromosomes with mapped rDNA SNPs (Lu et al. 2018) (see Reagent Table). Stocks used to study female nucleolar dominance were obtained from the UC San Diego *Drosophila* Stock Center and the culture was established by using single pair parents to minimize heterogeneity of rDNA within each stock.

The X and Y chromosomes from wild type (*yw*) were introduced to genotypes of interest analyzed in this study to keep their rDNA loci consistent across experiments. When it was not feasible to introduce wild type (*yw*) X/Y chromosomes into a genetic background of interest, their rDNA was sequenced to find SNPs between the X and Y rDNA (see below).

### RNA in situ hybridization

Third instar larval or adult tissues were dissected in RNase-free 1X PBS, fixed in RNas-free 4% formaldehyde, and incubated overnight in 70% EtOH at 4° to permeabilize the tissues. Embryos were collected according to a modified protocol from (Wilk et al. 2010) by allowing parents to lay eggs on an apple-agar plate at room temperature for a range of collection time (3 – 17hrs). Embryos were transferred to glass scintillation vials with glass Pasteur pipettes and were washed of any yeast in 1X PBS then dechorionated in 50% bleach for 30 sec and washed again in PBS. The embryos were then devitellinized and fixed in 50:50 heptane:4% RNas-free formaldehyde during vigorous, manual shaking for 20 min, then again in 50:50 heptane:methanol twice for 30 sec, washed in methanol then stored in methanol at −20° for at least one night before proceeding to *in situ* hybridization.

*In situ* hybridization was performed as previously described with slight modifications (Lu et al. 2018). In short, samples were washed with wash buffer (10% formamide in 1X SSC and 0.1% Tween-20) for 5 min, then incubated with the hybridization mix (10% formamide, 1X SSC, 10% Dextran sulfate (w/v) (Sigma, D8906), 100nM each *in situ* probe, and 300nM each mask oligo (for SNP *in situ*) overnight in a 37° water bath. Samples were then washed twice in wash buffer for 30 min each at 37° and stored in Vectashield H-1200 (Vector Laboratories) with DAPI. Hybridization and washes for X/X females were performed at 42°. Images were taken using a Leica TCS SP8 confocal microscope with 63X oil-immersion objectives and processed using Adobe Photoshop software.

See Reagent Table for fluorescent *in situ* oligonucleotide probes. SNP *in situ* oligonucleotide probes were custom ordered from Biosearch Technologies (Lu et al. 2018). Fluorescent *in situ* oligonucleotide probes used to study female nucleolar dominance were designed using Integrated DNA Technologies Oligo Analyzer.

### Identification of SNPs in rDNA

To sequence X rDNA, genomic DNA was extracted from 10-15 female flies of a genotype of interest. To sequence Y rDNA, male flies of the genotype of interest were crossed to C(1)DX/Y female flies, which lack X rDNA, and 10-15 female progeny (which have the Y chromosome of interest and C(1)DX) was subjected to genomic DNA extraction. PCR was performed on the extracted genomic DNA to amplify three regions of the rDNA with the following primers:

18S: (forward) 5’-GAAACGGCTACCACATCTAAGG-3’ and (reverse) 5’-GGACCTCTCGGTCTAGGAAATA-3’.

ITS1: (forward) 5’-CTTGCGTGTTACGGTTGTTTC-3’ and (reverse) 5’-ACAGCATGGACTGCGATATG-3’.

28S: (forward) 5’-AGCCCGATGAACCTGAATATC-3’ and (reverse) 5’-CATGCTCTTCTAGCCCATCTAC-3’ (Lu et al. 2018).

PCR products were verified by agarose gel electrophoresis and purified using a PCR Purification Kit (Qiagen). Sanger sequencing was performed on the purified PCR products using the same PCR primers (University of Michigan Biomedical Research DNA Sequencing Core Facility). Sequencing data was analyzed using the free downloadable software ApE: A plasmid Editor, by M. Wayne Davis. 4 SNPs between the X and Y rDNA were previously found (Lu et al. 2018) and a combination of 3-4 of the SNP probes were used depending on the presence or absence of SNPs in the specific strain (see Reagent Table). Unless otherwise stated, all 4 SNP probes (SNPs 1-4) were used for *in situ* hybridization.

### Larval brain squash and DNA FISH on mitotic chromosomes

We utilized a modified DNA fluorescent *in situ* hybridization (FISH) protocol described previously (Larracuente and Ferree 2015, Jagannathan et al. 2017). In short, third instar larvae were collected and brains were dissected in 1X PBS. Larval brains were fixed in 45:55 acetic acid:4% formaldehyde in PBS on Superfrost Plus Microscope Slides (Fisherbrand 22-037-246). The sample was then covered with a coverslip, manually squashed, and submerged in liquid nitrogen until frozen. The coverslips were quickly removed and the slides were treated with 100% ethanol at room temperature for 5 min. 20μL of hybridization buffer (50% formamide, 10% dextran sulfate, 2X SSC buffer, 0.5μM of each probe) was added to the sample, covered with a cover slip, and the sample was heat-denatured at 95° for 5 min, followed by incubation in a humid chamber in the dark overnight at room temperature. Samples were washed three times for 15 min in 0.1X SSC and then mounted in Vectashield H-1200 (Vector Laboratories) with DAPI. Probe sequences are provided in the Reagent Table.

### Immunofluorescence on mitotic chromosome spreads

A protocol described in (Blum et al. 2017) was used to conduct immunofluorescence on mitotic chromosome spreads. Briefly, larval brains from third instar larvae were dissected and incubated in 30μL of 0.5% sodium citrate on Superfrost Plus Microscope Slides (Fisherbrand 22-037-246) for 10-20 min. Sodium citrate was gently removed using a micropipette. 25μL of 4% formaldehyde was gently added to the slide over the sample, removed with a micropipette and replaced with another fresh 25μL of 4% formaldehyde and fixed for 4 min. During fixation, the larval brains were dissected into smaller pieces. Any imaginal discs and/or the ventral nerve cord were removed during this process. After fixation, the sample was covered with a coverslip, squashed and submerged in liquid nitrogen until frozen. After removal of the coverslips, slides were washed in PBS for 30 - 60 min and incubated overnight with primary antibodies (Chicken anti-Cid, 1:200) in 3% BSA in 1X PBST at 4° in a humid chamber. The slides were washed in 1X PBST, three times for 20 min each, then incubated with secondary antibodies (Goat anti-Chicken Alexa Fluor 488 Invitrogen, A-11039, 1:200) in 3% BSA in 1X PBST for 45 min at room temperature in a humid chamber in the dark. Slides were washed in 1X PBST, three times for 20 min and mounted in Vectashield H-1200 (Vector Laboratories) with DAPI. Antibodies are listed in the Reagent Table.

### Quantification and statistical analysis of nucleolar dominance

Nucleolar dominance was quantified manually from images generated using a Leica TCS SP8 confocal microscope. For each embryo, larval brain, imaginal disc, larval anterior midgut, and adult anterior midgut sample, 1-3 representative images of each tissue were captured for scoring purposes. Imaginal discs were randomly scored without intentionally excluding any imaginal disc type, therefore all imaginal discs were included in the category of ‘discs’ for scoring purposes. Z-stacks were generated with maximum projections for pre-gastrulation embryos, larval anterior midgut and adult anterior midgut images for scoring. Whole tissues were scored for salivary glands and larval fat bodies. All cells were identified and scored based on nuclear DAPI staining and morphology. Note that call of dominance vs. co-dominance was straightforward, owing to consistent signal intensity across samples based on the RNA *in situ* procedure described above. The number of cells and the number of tissues scored per genotype are listed in each corresponding figure legend and in Table S1. p-values were calculated using Welch’s unpaired, unequal variances Student’s t-test with n representing number of tissues scored.

### Data availability

*Drosophila* strains and reagents are listed in the Reagent Table and/or above. Raw scoring data are provided in Table S1.

## Results

### Y rDNA dominance is gradually established during male development

Thorough characterization of nucleolar dominance within a species (i.e. in the context of non-hybrids) has been limited to *A. thaliana* (Tucker et al. 2010). To extend the analysis of nucleolar dominance in *D. melanogaster*, which has been limited to larval neuroblasts and adult male germline cells (Greil and Ahmad 2012, Lu et al. 2018), we applied the SNP *in situ* hybridization method that differentiates X rDNA-derived rRNA vs. Y rDNA-derived rRNA (Lu et al. 2018) and comprehensively analyzed the state of nucleolar dominance during development of *D. melanogaster* (Figure 1A). In all experiments reported in this study, X and Y chromosomes with defined rDNA SNPs from a wild type strain (*yw*) were introduced into the genetic background of interest. Alternatively, distinct SNPs were identified by sequencing X and Y rDNA loci, if introduction of the *yw* strain sex chromosomes was complicated/impossible (see Materials and Methods).

**Figure 1:**
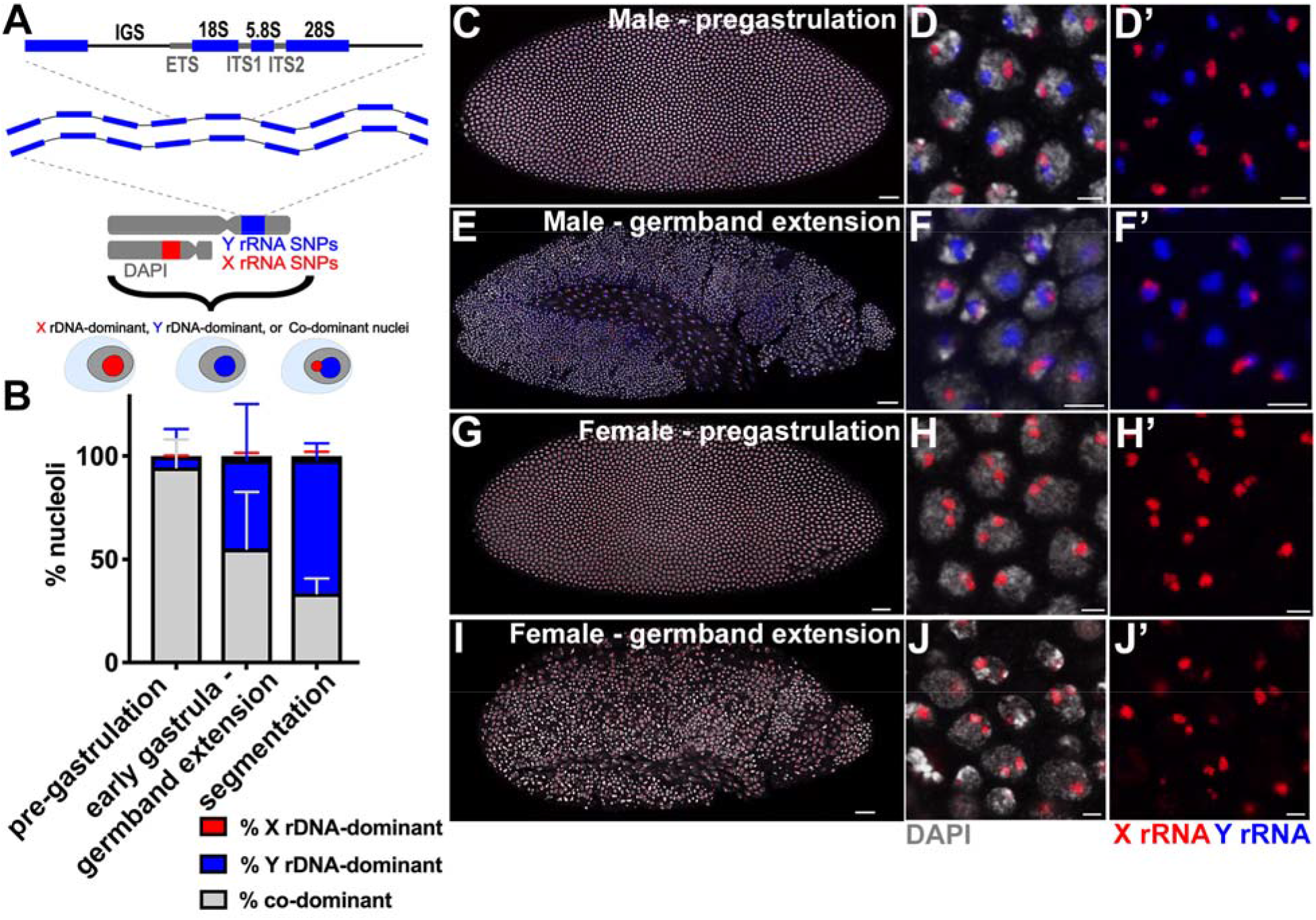
Nucleolar dominance is not established during embryogenesis in male embryos. A) Schematic of rDNA repeats (IGS = intergenic spacer, ETS = external transcribed spacer, ITS = internal transcribed spacer, 18S, 5.8S and 28S = rRNA coding region). X and Y rDNA can be distinguished by SNP *in situ* hybridization. Definition of X rDNA-dominant, Y rDNA-dominant, or co-dominant is shown. B) Quantification of nucleolar dominance across embryogenesis in males: pre-gastrulation (n = 748 cells from 7 embryos), early gastrula through germband extension (n = 1086 cells from 12 embryos), and segmentation (n = 1242 cells from 10 embryos). Red = % X rDNA-dominant, blue = % Y rDNA-dominant, grey = % co-dominant. C) Male pre-gastrulation embryo, scale = 25μm, D) zoomed image of nuclei from male pre-gastrulation embryo, scale = 3μm. E) Male embryo at germband extension stage, scale = 25μm, F) zoomed image of male embryo at germband extension stage, scale = 3μm. G) Female pre-gastrulation embryo, scale = 25μm, H) zoomed image of female pre-gastrulation embryo, scale = 3μm. I) Female embryo at germband extension stage, scale = 25μm, J) zoomed image of female embryo at germband extension stage, scale = 3μm. Red = X rRNA, blue Y rRNA, white = DAPI.

We first focused on nucleolar dominance in male embryos: 48.6% of the total embryos scored (n = 368) contained both the Y rDNA and X rDNA SNP signals, which we deemed as male embryos. Note that not all nuclei within an embryo necessarily contained both X rRNA and Y rRNA signals, but the presence of any Y rRNA-containing nuclei within an embryo indicates that they are male embryos. On the contrary, 51.4% of embryos contained only X rDNA SNP signal in all nuclei within an entire embryo, which were deemed as female embryos. Since our SNP *in situ* probes cannot discriminate rRNA signals from two X rDNA loci in females, their state of nucleolar dominance cannot be determined by these experiments (Figure 1G-J) (see below for nucleolar dominance in females). We found that in early male embryos (pre-gastrulating, around syncytial cycle 13-14), the majority of nuclei expressed both the X and Y rDNA (i.e. co-dominant) (94.8 ± 13.2%) (Figure 1B-D). It has been reported that larval neuroblasts (Greil and Ahmad 2012), male germline stem cells (GSCs) and spermatogonia (Lu et al. 2018) exhibit Y rDNA dominance, suggesting that nucleolar dominance may be established during the course of development. To address this possibility, we examined the state of nucleolar dominance along the course of development through embryonic stages, larval development and into adulthood. Although the pre-gastrulating embryos exhibited high frequency of co-dominance (∼95%), we observed a decrease in the percentage of co-dominant nuclei, with a concomitant increase in Y rDNA-dominant cells as male embryos progressed through development (Figure 1B-F). Male embryos during early gastrula or germband extension stages show 55.3 ± 27.5% co-dominant nuclei and 42.9 ± 27.0% Y rDNA dominant nuclei (Figure 1B-F). Later during segmentation, co-dominant nuclei further decreased to 33.7 ± 7.1%, as Y rDNA dominant nuclei increased to 64.6 ± 7.9% (Figure 1B).

As development proceeds to the larval stage, we observed much higher rates of Y rDNA dominance in most tissues: larval brains (83.5 ± 4.6%) similar to what has been previously reported (Greil and Ahmad 2012), imaginal discs (93.6 ± 3.1%), larval fat bodies (95.8 ± 5.1%), and larval anterior midgut enterocytes (82.9 ± 12.1%) (Figure 2A, 2C-I). Salivary glands, which undergo a high degree of polyploidization, showed only a moderate degree of Y rDNA dominance (51 ± 19.4%) (Figure 2A-B). Y rDNA dominance in the anterior midgut further increased in the adult (from 82.9% ± 12.1% in third instar larvae to 99.7 ± 0.7% in adult) (Figure 2A, 2J-K). These data suggest that nucleolar dominance in *D. melanogaster* males is gradually established over the course of development. This is similar to what was reported in *Arabidopsis* (Pontes et al. 2007, Earley et al. 2010), where seedling cotyledons exhibit co-dominance and nucleolar dominance is established in later stages of development.

**Figure 2:**
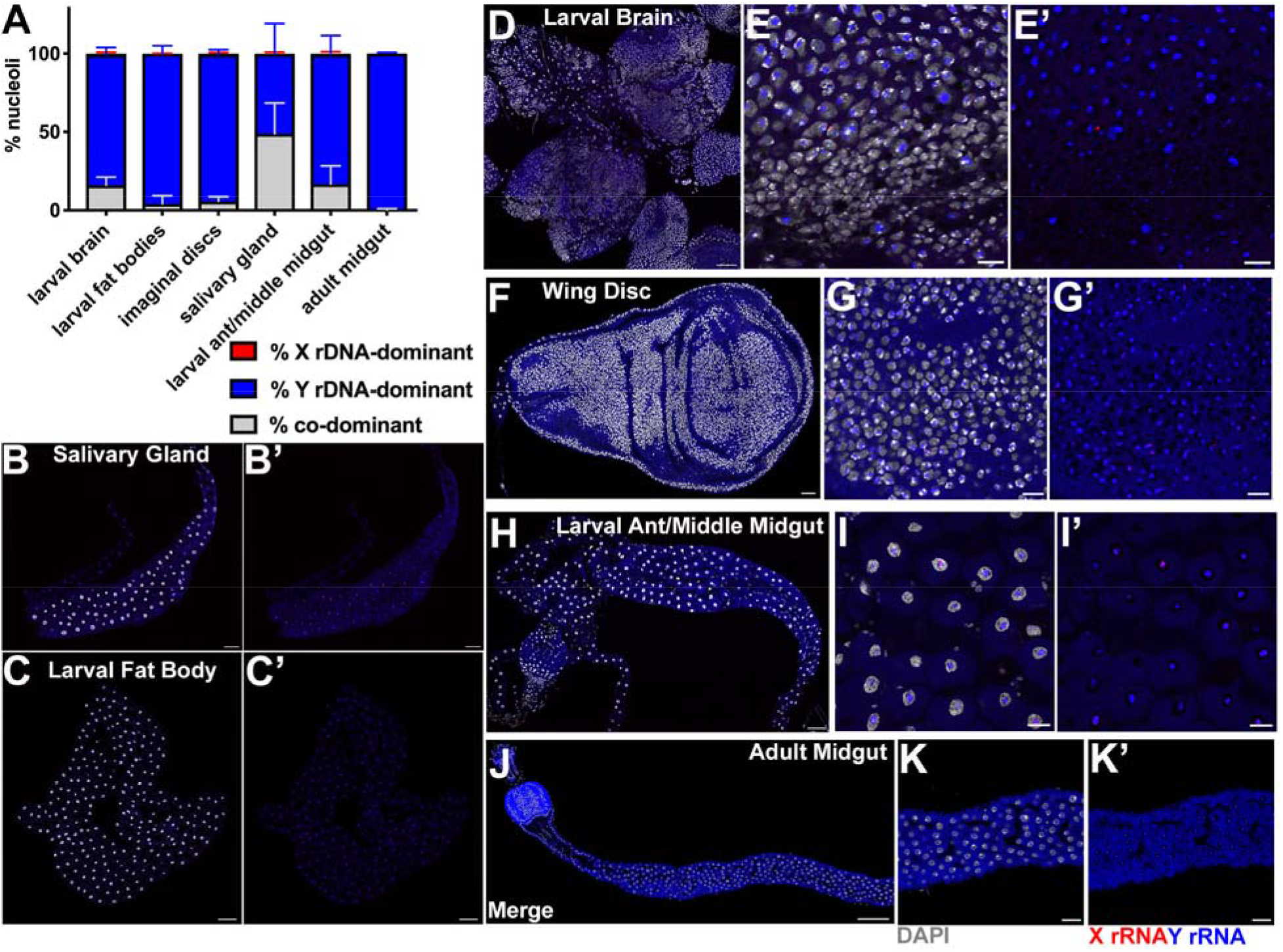
Y rDNA dominance is established during development in males. A) Quantification of nucleolar dominance in larval and adult tissue(s) in males: larval brain (n = 1594 cells from 6 brains), larval fat bodies (n = 1575 cells from 17 fat bodies), imaginal discs (n = 1251 cells from 5 imaginal discs), salivary gland (n = 878 cells from 15 salivary glands), larval anterior midgut (n = 81 cells from 6 guts), adult anterior midgut (n = 922 cells from 7 guts). Red = % X rDNA-dominant, blue = % Y rDNA-dominant, grey = % co-dominant. B) Representative images of whole mount salivary gland, scale = 100μm, C) larval fat body, scale = 100μm, D) larval brain, scale = 50μm, E) zoomed image of larval brain, scale = 10μm, F) wing disc, scale = 25μm, G) zoomed image of wing disc, scale = 8μm, H) larval anterior midgut, scale = 100μm, l) zoomed image of larval anterior midgut, scale = 25μm, J) adult anterior midgut, scale = 100μm, K) zoomed image of adult anterior midgut, scale = 25μm. Red = X rRNA, blue Y rRNA, white = DAPI.

### Histone methyltransferase Su(var)3-9 aids in the establishment of Y rDNA dominance in males across tissues

Small interfering RNAs (siRNAs) in *Arabidopsis* (Preuss et al. 2008) and long non-coding, promoter-associated RNAs in mammalian cell lines (Mayer et al. 2006) were shown to regulate rDNA silencing. These non-coding RNAs recruit factors that induce heterochromatinization of rDNA through DNA methylation (Lawrence et al. 2004, Preuss et al. 2008, Schmitz et al. 2010), histone methylation (Lawrence et al. 2004, Santoro and Grummt 2005, Pontvianne et al. 2012) and histone deacetylation (Santoro and Grummt 2005, Earley et al. 2006). The small RNA pathway and heterochromatin factors have also been shown to influence nucleolar morphology in *D. melanogaster* larval tissues, which may reflect disrupted rDNA expression (Peng and Karpen 2007). Based on these previous studies, we wondered whether the small RNA machinery and/or heterochromatin formation play a role in nucleolar dominance in *D. melanogaster*. To test this, we assessed nucleolar dominance in the mutants of *dicer-2*, an endonuclease critical for the siRNA pathways, or *Su(var)3-9*, a histone methyltransferase critical for depositing heterochromatin-associated histone methylation. We found that *dcr-2^L811fsx^/dcr-2^p[f06544]^* mutants showed only a slight (although statistically significant) change in Y rDNA dominance in larval brains (71.4 ± 7.5% compared to control 83.5 ± 4.6%) and imaginal discs (81.8 ± 10.9% compared to control 93.6 ± 3.1%) (Figure 3A-B, 3C-D, 3F-G), suggesting that the siRNA mechanism might not play an important role in nucleolar dominance, as opposed to what is reported in *Arabidopsis* (Pontes et al. 2006, Preuss et al. 2008). *Su(var)3-9^1^/Su(var)3-9^2^* mutants showed a marked decrease in Y rDNA dominance in larval brain (46.5 ± 16.5%, compared to control 83.5 ± 4.6%) and imaginal discs (61.4 ± 21.0%, compared to control 93.6 ± 3.1%) (Figure 3A-B, 3C, 3E, 3F, 3H), consistent with the previous finding that *Su(var)3-9* is involved in silencing of X rDNA in male neuroblasts (Greil and Ahmad 2012). *dicer-2* mutants and *Su(var)3-9* mutants had minimal effects on nucleolar dominance in polyploid tissues (salivary glands, larval fat bodies, larval anterior midgut, and adult anterior midgut enterocytes) (Figure S1A-D). These results suggest that the small RNA pathway is most likely not important for Y rDNA dominance, whereas heterochromatin formation is important for Y rDNA dominance in diploid tissues in *D. melanogaster*.

**Figure 3:**
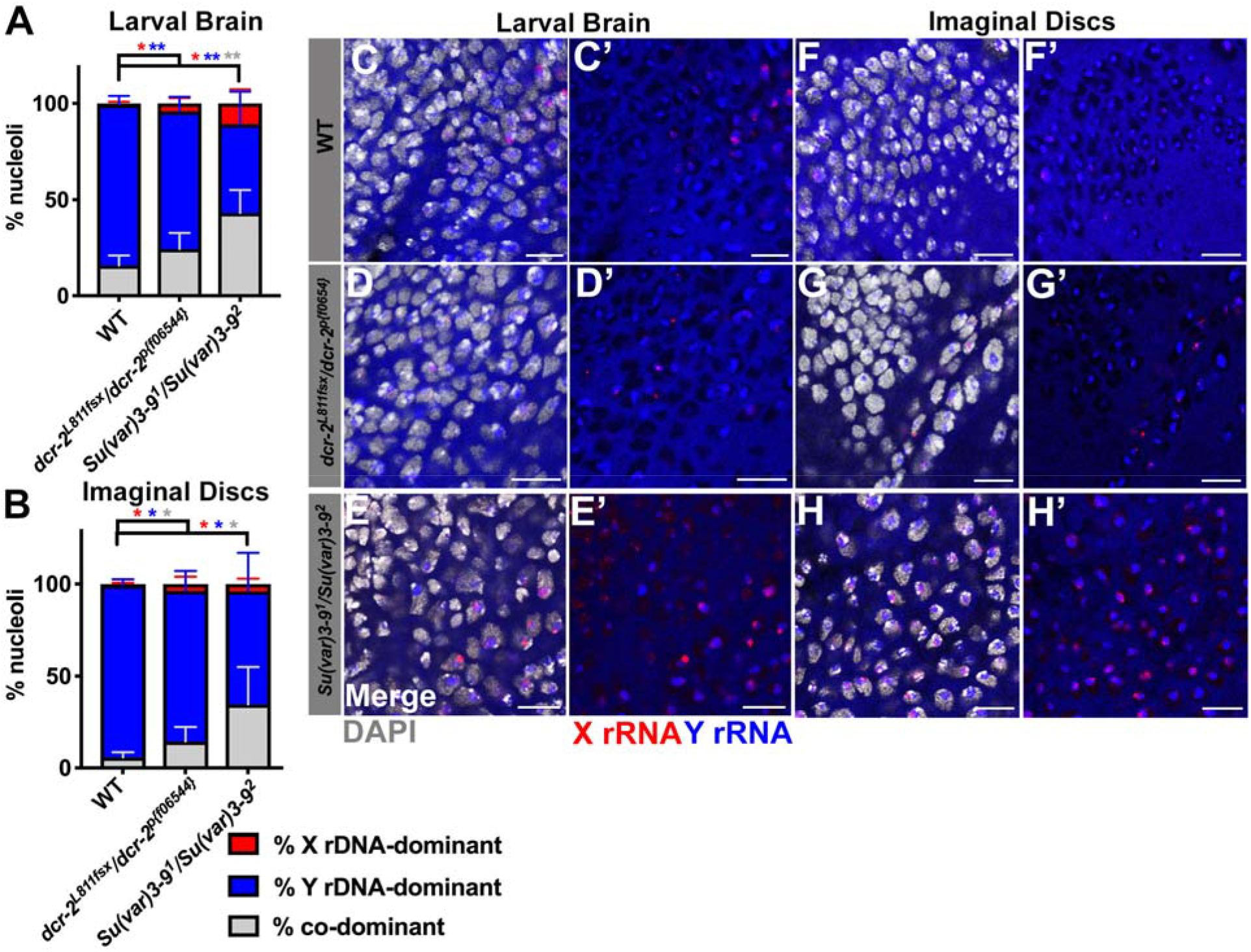
Heterochromatin formation aids in Y rDNA dominance in males. Quantification of male nucleolar dominance in A) larval brains of wild type (*yw*) (n = 1594 cells from 6 brains), *dcr-2^L811fsx^/dcr dcr-2^p[f06544]^* mutants (n = 504 cells from 6 brains), and *Su(var)3-9^1^/Su(var)3-9^2^* mutants (n = 461 cells from 6 brains) (the same wild type data from Figure 2 for comparison). B) Quantification of nucleolar dominance in male imaginal discs of wild type (n = 1251 cells from 5 imaginal discs), *dcr-2^L811fsx^/ dcr-2^p[f06544]^* mutants (n = 579 cells from 9 imaginal discs), and *Su(var)3-9^1^/Su(var)3-9^2^* mutants (n = 432 cells from 6 imaginal discs) (the same wild type data from Figure 2 for comparison). Red = % X rDNA-dominant, blue = % Y rDNA-dominant, grey = % co-dominant nuclei. p-values calculated using Welch’s unpaired, unequal variances t-test using n = number of tissues. (no star) = not significant, * = < 0.05, ** = < 0.01. Colors of asterisks correspond to colors of bars for which P-values were calculated (e.g. blue asterisk for Y rDNA-dominant p-values). Representative images of larval brains from C) wild type (*yw*), D) *dcr-2^L811fsx^/ dcr-2^p[f06544]^* mutants, and E) *Su(var)3-9^1^/Su(var)3-9^2^* mutants. Representative images of imaginal discs from F) wild type, G) *dcr-2^L811fsx^/ dcr-2^p[f06544]^*mutants, and H) *Su(var)3-9^1^/Su(var)3-9^2^* mutants. Red = X rRNA, blue Y rRNA, white = DAPI. All scale bars = 10μm.

### Nucleolar dominance does not occur in X/X females across tissues

It was previously shown that nucleolar dominance does not occur in female larval neuroblasts (Greil and Ahmad 2012). We sought to determine the state of nucleolar dominance in females (between two X rDNA loci) across tissues and developmental stages. Doing so requires two distinct X rDNA loci with detectable differences, similar to SNP *in situ* hybridization described above for X vs. Y rDNA. Our initial searches for SNPs between X rDNA loci from multiple laboratory strains revealed no SNPs (see Materials and Methods). However, sequencing of X rDNA from geographically separated *D. melanogaster* strains led us to the identification of a 24-bp deletion in the internal transcribed spacer (ITS1) of the X rDNA in a strain originating from Guam, compared to most other strains sequenced (i.e. *yw*, Oregon-R, Canton-S, Beijing, Pohnpei, Samoa, Port Moresby, Le Réduit) (see Materials and Methods, Figure 4A). We designed oligonucleotide probes to distinguish the rRNA from the Guam strain (ITS^Δ24^) vs. other strains (ITS^+^) (see Materials and Methods, Figure 4A). The Guam strain exhibited signals only from ITS^Δ24^ (Figure S2). Among other strains that have the ITS^+^ variant, the Le Réduit strain had the least background signal with the ITS^Δ24^ probe (Figure S2), whereas females from other strains revealed weak ITS^Δ24^ signal in addition to predominant ITS^+^ signal (data not shown), possibly because these strains may contain a small fraction of rDNA copies with the ITS^Δ24^ variant. Based on these results, we decided to utilize the Guam and Le Réduit strains to determine the state of nucleolar dominance between two X rDNA loci in females.

**Figure 4:**
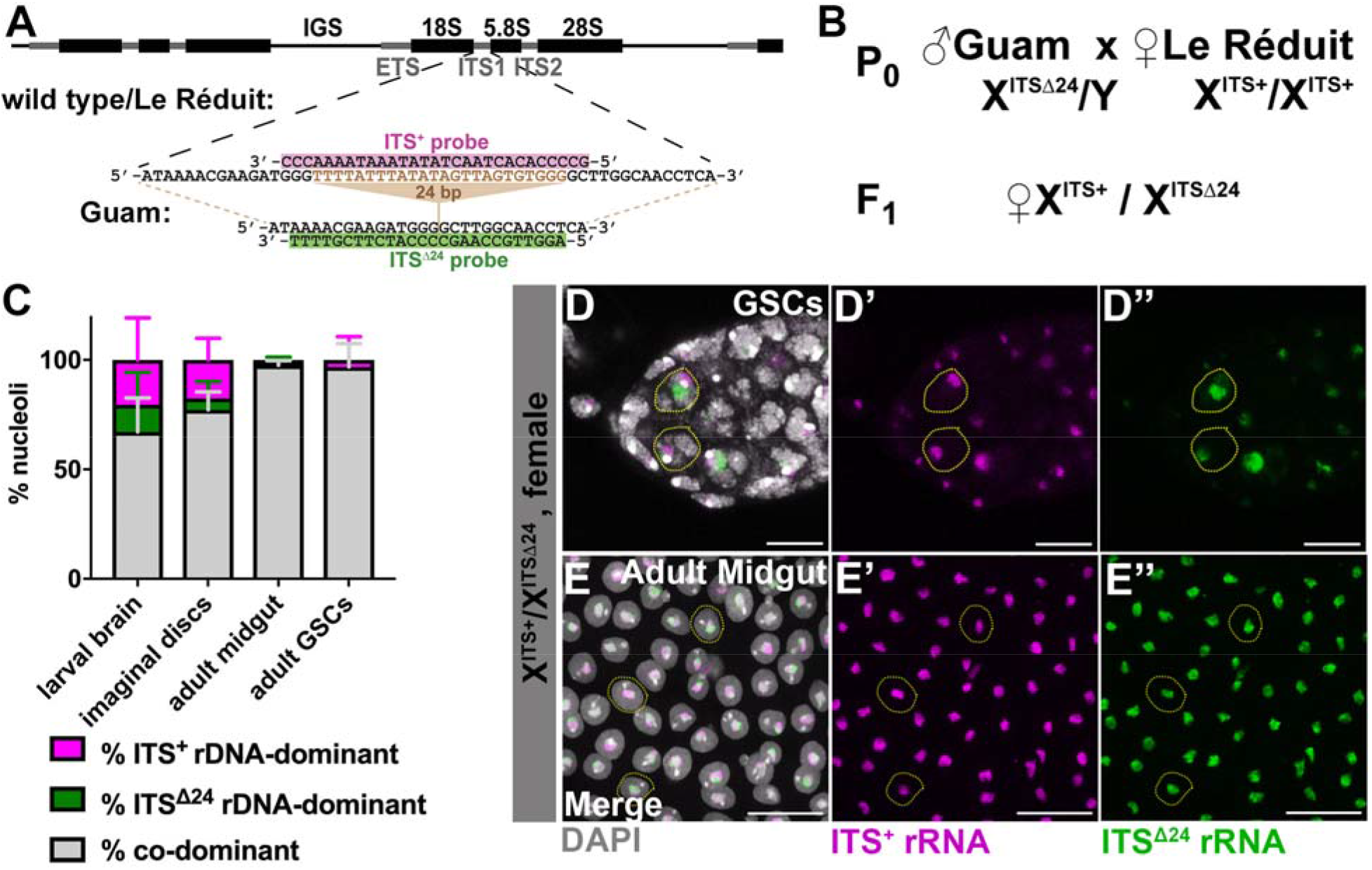
X/X females express both rDNA loci throughout development. A) Oligonucleotide probe design to differentially visualize two distinct X-chromosome rDNA internal transcribed spacer (ITS) transcripts, utilizing a 24-bp deletion in the rDNA ITS between wild type/ Le Réduit and Guam *D. melanogaster* strains. B) The cross to generate female F1 with one X chromosome with wild type ITS (ITS^+^) and the other X chromosome with the ITS with 24bp deletion (ITS^Δ24^). C) Quantification of nucleolar dominance between two X rDNA in female larval and adult tissue(s): larval brain (n = 2616 cells from 9 brains), imaginal discs (n = 2575 cells from 9 imaginal discs), adult anterior midgut (n = 904 cells from 9 guts), adult GSCs (n = 150 cells from 57 germarium). Magenta = % ITS^+^ rDNA-dominant, green = % ITS^Δ24^ rDNA-dominant, grey = % co-dominant nuclei. Representative images of D-D’’) GSCs, scale = 8μm, and E-E’’) adult anterior midgut enterocytes, scale = 25μm. Magenta = ITS^+^ rRNA, green = ITS^Δ24^ rRNA, white = DAPI.

We crossed Guam strain males with Le Réduit strain females and tissues from the resulting F1 females were assessed for the state of nucleolar dominance by RNA *in situ* using the ITS^Δ24^ and ITS^+^ probes (Figure 4B). We found that X/X female cells predominantly show expression from both rDNA loci (i.e. co-dominant) in larval brains (67.1 ± 15.6% co-dominant) (Figure 4C). We found that X/X female larval imaginal discs also exhibit primarily co-dominance (77.3 ± 8.1%) (Figure 4C). Adult tissues revealed even higher rates of co-dominance compared to larval tissues: anterior midgut enterocytes (97.6 ± 2.1% co-dominance) and GSCs (96.8 ± 10.7% co-dominance) (Figure 4C-E). It should be noted that we did not assess nucleolar dominance in female embryos because the Guam strain Y rDNA shares the same ITS sequence as Le Réduit X rDNA (ITS^+^), making the accurate sexing of embryos impossible. However, female embryos in the experiments described in Figure 1H-J mostly exhibited two nucleoli per nucleus, suggesting that female embryos also exhibit co-dominance.

The co-dominant state of X rDNA loci in the progeny of Guam and Le Réduit parents did not change even when parental origin of ITS^Δ24^ vs. ITS^+^ rDNA loci was switched (i.e. Guam females crossed to Le Réduit males) (Figure S3B). It should be noted that in this direction of the cross, females exhibited hybrid dysgenesis, leading to a high frequency of degenerated ovaries (Figure S3A). Despite high rates of hybrid dysgenesis, all cells scored exhibited co-dominance (Figure S3B). These results establish that females exhibit co-dominance in a broad range of tissues and developmental stages, extending the previous findings in female neuroblasts (Greil and Ahmad 2012).

### Y rDNA can dominate in female cells

The above results reveal a striking difference in the state of nucleolar dominance between males and females: Y rDNA dominates over X rDNA in males, whereas two X chromosomes are co-dominant in females. What differences between the X and Y chromosome determine the decision of which rDNA locus is to be expressed? A previous study showed that nucleolar dominance in *D. melanogaster* is not likely due to imprinting during the parents’ gametogenesis where the inheritance of the X and Y chromosomes is reversed (Greil and Ahmad 2012). Others have speculated that distinct sequence differences between the loci, in this case the X rDNA vs. Y rDNA loci, allow selective expression of particular rDNA loci (Kidd and Glover 1981, Macleod and Bird 1982, Labhart and Reeder 1984, Grimaldi et al. 1990, Heix and Grummt 1995, Neves et al. 1995, Houchins et al. 1997, Caudy and Pikaard 2002, Felle et al. 2010). Yet another possibility is that chromosomal context, or location within a particular chromosome (Chandrasekhara et al. 2016, Mohannath et al. 2016) may determine whether or not a particular rDNA locus may be expressed/silenced. In addition, cellular sex might determine whether or not nucleolar dominance occurs.

Because parental imprinting unlikely contributes to the regulation of nucleolar dominance (Greil and Ahmad 2012), we sought to test the possibility that X and/or Y rDNA contain specific *(cis)* elements that determine the state of nucleolar dominance. To this end, we examined the state of nucleolar dominance in females that carry a Y chromosome. C(1)RM is a compound X chromosome (two X chromosomes are fused and it contains one rDNA locus) and C(1)RM/Y flies develop as females (Bridges 1916, Bridges 1921). The rDNA on C(1)RM was found to share all SNPs with the *yw* X rDNA (see Materials and Methods and the Reagent Table). Utilizing these SNPs, we determined the state of nucleolar dominance between C(1)RM rDNA and Y rDNA in female tissues (e.g. diploid larval tissues, the adult anterior midgut and adult ovary) (Figure 5A-C, 5E). Surprisingly, C(1)RM/Y females exhibited strikingly high frequency of Y rDNA dominance in many cell types (Figure 5A-C, 5E). This is a stark contrast to X/X females, where two X chromosomes exhibit co-dominance across tissues (Figure 5A-C). These results indicate that Y rDNA dominance is determined by the Y chromosome (e.g. sequence information within Y rDNA or other elements on the Y chromosome), and disfavors the possibility that cellular sex determines dominance vs. co-dominance.

**Figure 5:**
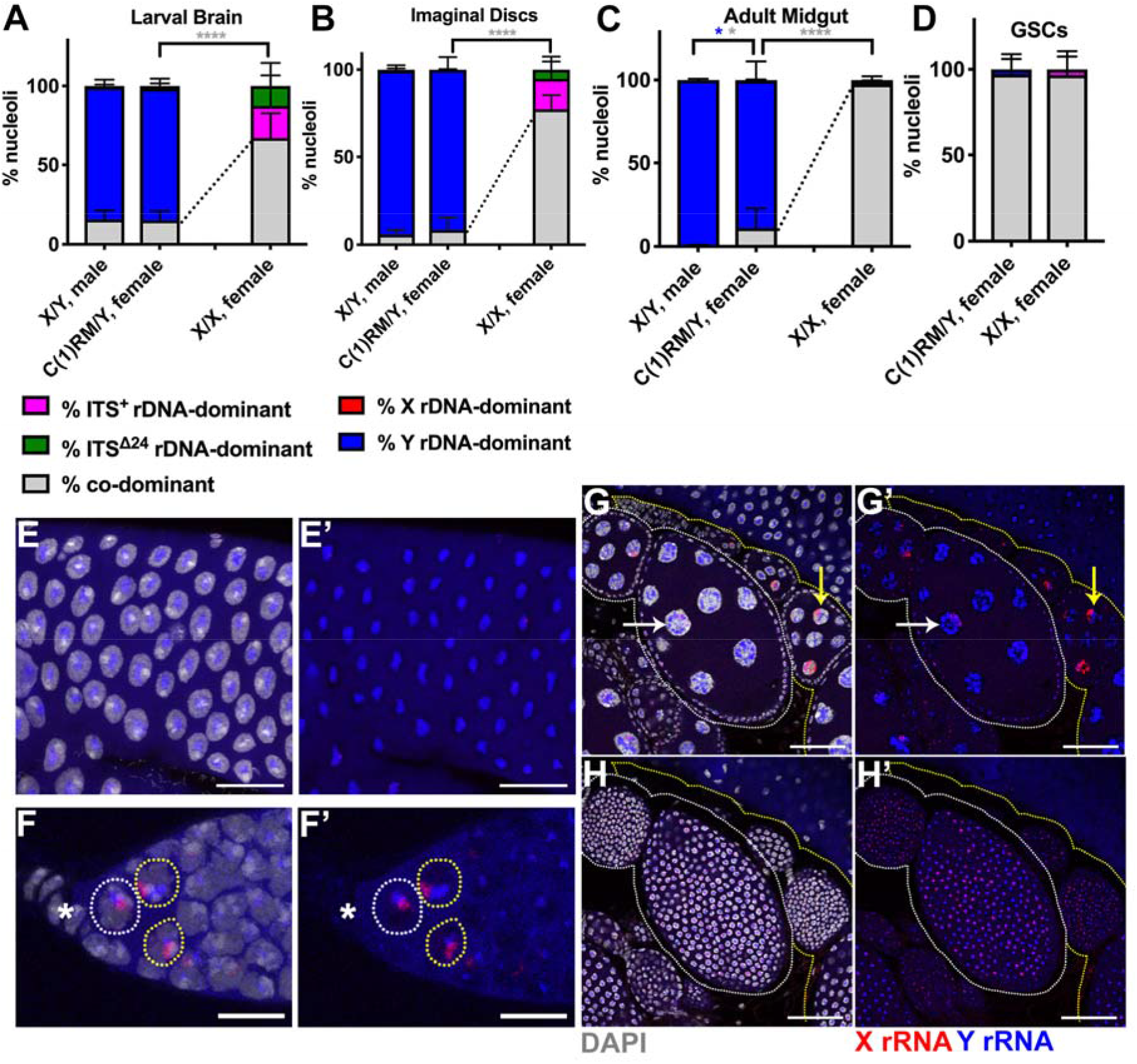
The Y rDNA can dominate over X rDNA in female cells. A-D) Quantification of nucleolar dominance in C(1)RM/Y females compared to males (data from Figure 2 for comparison) and typical X/X females (data from Figure 4 for comparison) in larval brain (A, n = 914 cells from 9 brains), imaginal discs (B, n = 1068 cells from 12 imaginal discs), adult anterior midgut enterocytes (C, n = 870 cells from 7 guts), and female GSCs (D, n = 54 cells from 21 germarium). Dotted lines denote differences in rates of co-dominance between C(1)RM/Y and typical X/X females. p-values calculated using Welch’s unpaired, unequal variances *t*-test using number of tissues scored. p-values between C(1)RM/Y and XX females were only calculated for % co-dominant. (no star) = not significant, * = < 0.05, **** = < 0.0001. Colors of asterisks correspond to colors of bars for which P-values were calculated (e.g. blue asterisk for Y rDNA-dominant p-values). E) Representative image of C(1)RM/Y female adult anterior midgut enterocytes, scale = 25μm. F) C(1)RM/Y female GSCs (white circle) and cystoblasts (yellow circles), * = terminal filament, scale = 10μm. G) two C(1)RM/Y ovarioles (separately circled in white or yellow), scale = 50μm. Arrows indicating nurse cells with low X rDNA expression (white) and high X rDNA expression (yellow). H) Follicle cells from C(1)RM/Y ovarioles corresponding to G (different Z-depth), scale = 50μm. Red = X rRNA, blue Y rRNA, white = DAPI.

Interestingly, adult female GSCs and cystoblasts showed a high degree of co-dominance in C(1)RM/Y females (Figure 5D, 5F), whereas some nurse cells showed Y rDNA dominance (Figure 5G). The somatic follicle cells of the egg chambers showed mostly co-dominance (Figure 5H). These data together suggest that, whereas the Y rDNA can dominate irrespective of cellular sex, it is not the sole factor to determine nucleolar dominance, and that cell-type specific information can modulate the state of nucleolar dominance.

### Y rDNA cis elements contribute to nucleolar dominance

The above data that the Y rDNA can dominate over the X rDNA irrespective of cellular sex in most cell types indicate that the Y chromosome may contain cis-acting element(s) that establish Y rDNA dominance. Such information may be embedded in the Y rDNA locus itself, such as variable sequences in the coding and/or spacer sequences (Tautz et al. 1987, Tautz et al. 1988, Schlotterer et al. 1994, Caudy and Pikaard 2002). Additionally, the entire chromosomal context may dictate the state of silencing/activation (Chandrasekhara et al. 2016, Mohannath et al. 2016). To address whether the Y rDNA contains *cis* information that influences its expression/dominance, we utilized an X chromosome that contains Y rDNA due to chromosomal rearrangements. In this chromosome, In(1)sc^4L^sc^8R^+Tp(1;YS)bb^AM7^ (referred as to X^bb−^ (Y^bb+^)), the original X rDNA locus was replaced with the rDNA locus from the Y chromosome (Figure 6A and Figure S4). We first sequenced the rDNA from this X^bb−^ (Y^bb+^) chromosome and found that its rDNA shared 3/4 of the SNPs with the *yw* Y rDNA (see Materials and Methods and the Reagent Table). Using these three SNP *in situ* probes, we found that X^bb−^(Y^bb+^)/X females exhibit intermediate patterns of nucleolar dominance: in larval brain, imaginal discs, and adult anterior midgut enterocytes, X^bb−^ (Y^bb+^) rDNA mostly dominates over X rDNA, as opposed to co-dominance in typical X/X females (Figure 6B-D, 6F, 6G, 6I-J). However, the degree of X^bb−^(Y^bb+^) rDNA dominance was lower than Y rDNA dominance in X/Y males (Figure 6B-C, 6J) and in C(1)RM/Y females (Figure 6D-E, 6G-H). GSCs from X^bb−^(Y^bb+^)/X females exhibited high rates of co-dominance, similar to X/X females (Figure 6K) as well as C(1)RM/Y females (compare to Figure 5D). These results suggest that Y rDNA carries critical information that allows for dominance of the Y rDNA locus. Additionally, the observation that the degree of Y rDNA dominance is much less than that in X/Y males or C(1)RM/Y females indicates that the chromosomal context (e.g. being embedded in the entire Y chromosome) also plays an important role in the determination of Y rDNA dominance.

**Figure 6:**
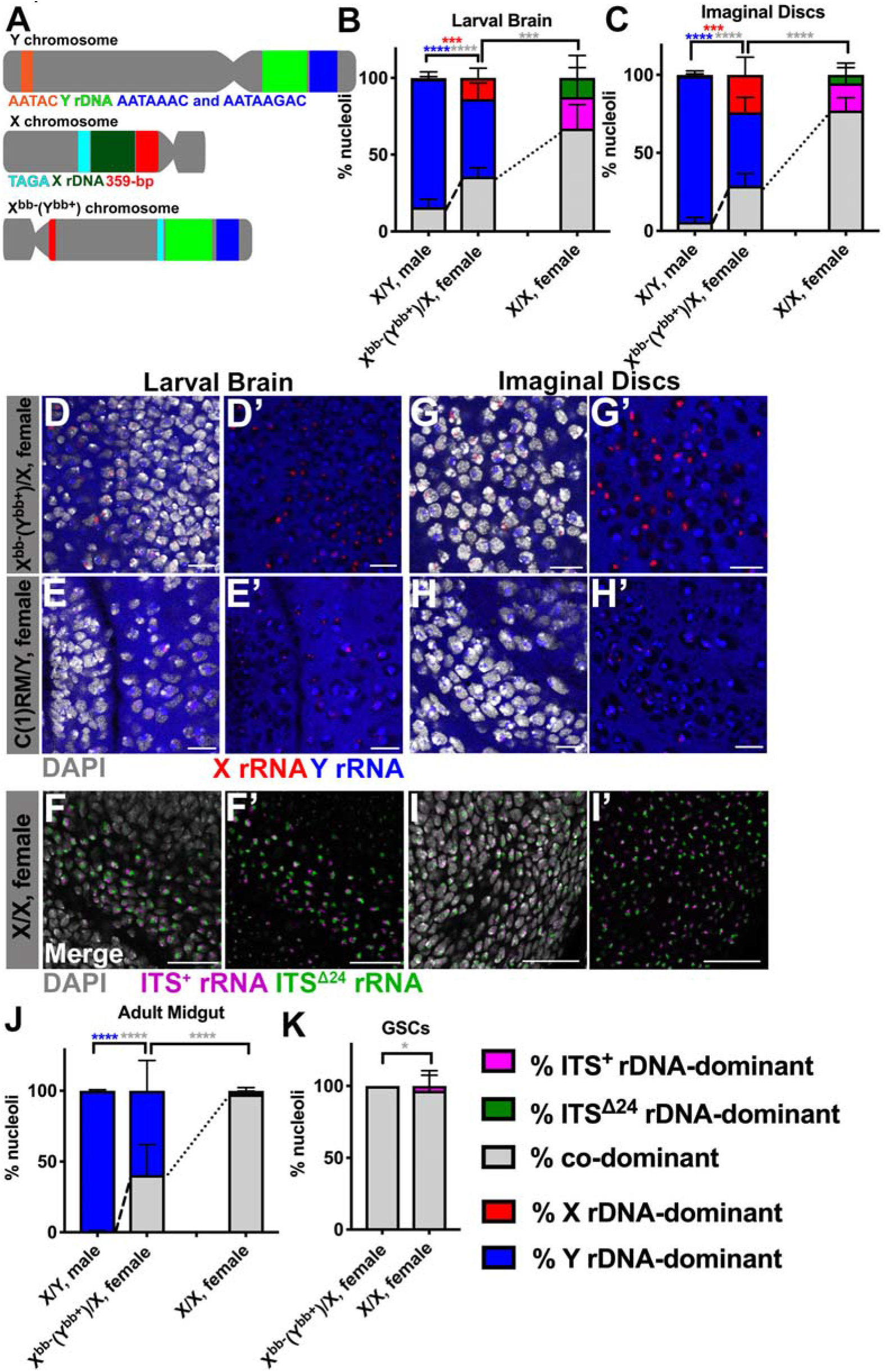
Y rDNA likely has *cis* information to dictate nucleolar dominance. A) Structure of a wild type Y chromosome, a wild type X chromosome, and the X^bb−^(Y^bb+^) chromosome based on Figure S4. Constriction represents centromere location. B-C) Quantification of nucleolar dominance in X^bb−^(Y^bb+^)/X females compared to X/Y males (data from Figure 2 for comparison) and X/X females (data from Figure 4 for comparison) in larval brain (B, n = 1400 cells from 8 brains) and imaginal discs (C, n = 1362 cells from 9 imaginal discs). Red = % X rDNA-dominant, blue = % Y rDNA-dominant, grey = % co-dominant, magenta = % ITS^+^ rDNA-dominant, green = % ITS^Δ24^ rDNA-dominant nuclei. p-values calculated using Welch’s unpaired, unequal variances t-test using n = number of tissues. p-values between X^bb−^(Y^bb^+)/X and XX females were only calculated for % co-dominance. Dotted lines denote differences in rates of co-dominance between X^bb−^(Y^bb+^)/X and X/X females. Dashed lines denote differences in rates of co-dominance between X^bb−^(Y^bb^+)/X females and X/Y males. *** = < 0.001, **** = < 0.0001. Colors of asterisks correspond to colors of bars for which P-values were calculated (e.g. blue asterisk for Y rDNA-dominant p-values). Representative images of larval brain from D) X^bb−^(Y^bb+^)/X females, scale = 10μm, E) C(1)RM/Y females, scale = 10μm, and F) X/X females, scale = 25μm. Representative images of imaginal discs from X^bb−^(Y^bb+^)/X females, scale = 8μm, H) C(1)RM/Y females, scale = 10μm, I) X/X females, scale = 25μm. Red = X rRNA, blue Y rRNA, white = DAPI, magenta = ITS^+^ rDNA transcript, green = ITS^Δ24^ rDNA transcript. J) Quantification of nucleolar dominance in X^bb−^(Y^bb+^)/X females compared to both X/Y males (data from Figure 2 for comparison) and X/X females (data from Figure 4 for comparison) in adult anterior midgut enterocytes (n = 1213 cells from 13 guts), and K) female GSCs (n = 122 cells from 51 germarium). Red = % X rDNA-dominant, blue = % Y rDNA-dominant, grey = % co-dominant, magenta = % ITS^+^ rDNA-dominant, green = % ITS^Δ24^ rDNA-dominant nuclei. p-values calculated using Welch’s unpaired, unequal variances t-test. p-values between X^bb−^(Y^bb+^)/X and XX females were only calculated for % co-dominance. Dotted lines denote differences in rates of co-dominance between X^bb−^(Y^bb+^)/X and X/X females. Dashed lines denote differences in rates of co-dominance between X^bb−^(Y^bb+^)/X females and X/Y males. (no star) = not significant, * = < 0.05, **** = < 0.0001.

## Discussion

In this study, we conducted a thorough characterization of nucleolar dominance in *D. melanogaster*. Our study extends the previous discovery in *D. melanogaster* male larval neuroblasts that nucleolar dominance occurs within a species (Greil and Ahmad 2012) to a broader range of tissues and developmental stages. Our study shows that nucleolar dominance is a developmentally regulated process, being established gradually during the course of development. This is reminiscent of what was seen in *Arabidopsis* (Pontes et al. 2007, Earley et al. 2010), and supports the notion that nucleolar dominance is not limited to interspecies hybrids.

Earlier studies (Lawrence et al. 2004, Santoro and Grummt 2005, Earley et al. 2006, Earley et al. 2010, Greil and Ahmad 2012, Pontvianne et al. 2012), confirmed here, revealed heterochromatin formation as a critical aspect of nucleolar dominance. However, this likely reflects the need of heterochromatinization to silence rDNA loci that were chosen to be silenced, but does not provide the mechanism of ‘choice’ that dictates which particular rDNA loci are to be silenced or activated. Studies in interspecies hybrids of *Brassica* suggest that the ‘choice’ mechanism dictates which rDNA loci are silenced, instead of which loci are expressed (Chen and Pikaard 1997). In contrast, our results rather suggest that the Y rDNA locus has the information that allows it to be dominantly expressed. In experiments described in this study, where the Y rDNA was introduced into the context of females (C(1)RM/Y and X^bb−^(Y^bb+^)/X), the Y rDNA exhibited a high degree of dominance even in female cells, suggesting that the Y rDNA harbors certain information that promotes its transcription. However, our data (Figure 5 and Figure 6) also reveals that additional factor(s) on the Y chromosome, not just its rDNA locus, are important for complete establishment of Y rDNA dominance. Our results, compared to the studies in *Arabidopsis* (Chen and Pikaard 1997), suggest that the ‘choice’ mechanism may vary across species.

Elements within rDNA have been shown to influence nucleolar dominance in interspecies hybrids, particularly the non-coding region of rDNA: the length of the intergenic spacer sequence (IGS), which contains rDNA promoters (Coen and Dover 1982), was shown to dictate dominance in interspecies *Xenopus* hybrids (Reeder et al. 1983). Because the IGS sequences are known to be highly diverged compared to the coding region of rDNA (Tautz et al. 1987, Tautz et al. 1988), it is tempting to speculate that differences in IGS sequences between X and Y rDNA loci dictate Y rDNA dominance. Indeed, differential activity of X rDNA IGS vs. Y rDNA IGS as promoter/enhancer for RNA polymerase I, suggested in a previous study (Labhart and Reeder 1984), may be an underlying mechanism for nucleolar dominance.

In summary, our work expands on previous studies in *Arabidopsis* and *Drosophila* and supports the notion that nucleolar dominance is not constrained to interspecies hybrids, and represents a mechanism of rRNA regulation within a species. Our study suggests that the Y rDNA may have *cis* elements that dictate Y rDNA’s dominance in *D. melanogaster*. The precise identity of the *cis* element(s) of the Y rDNA/Y chromosome, and how they mediate its preferential transcription await future investigation. Our study lays the foundation to identify *cis* elements that regulate nucleolar dominance and to understand the underlying mechanisms needed to achieve nucleolar dominance. Most importantly, why a locus-wide mechanism, i.e. nucleolar dominance, has evolved to regulate rDNA expression is a fundamental question to be addressed in the future.

## Acknowledgements

We thank Dr. Amanda Larracuente, Bloomington *Drosophila* Stock Center, Kyoto *Drosophila* Stock Center, and the UC San Diego *Drosophila* Stock Center for reagents. We thank the Yamashita lab members and Dr. Sue Hammoud for discussion and/or comments on this manuscript. This research was supported by Howard Hughes Medical Institute (to Y.Y) and in part by the NIH Career Training in Reproductive Biology (5T32HD079342-04) (to N.W-P).

## Supplementary Figures

**Figure S1:**
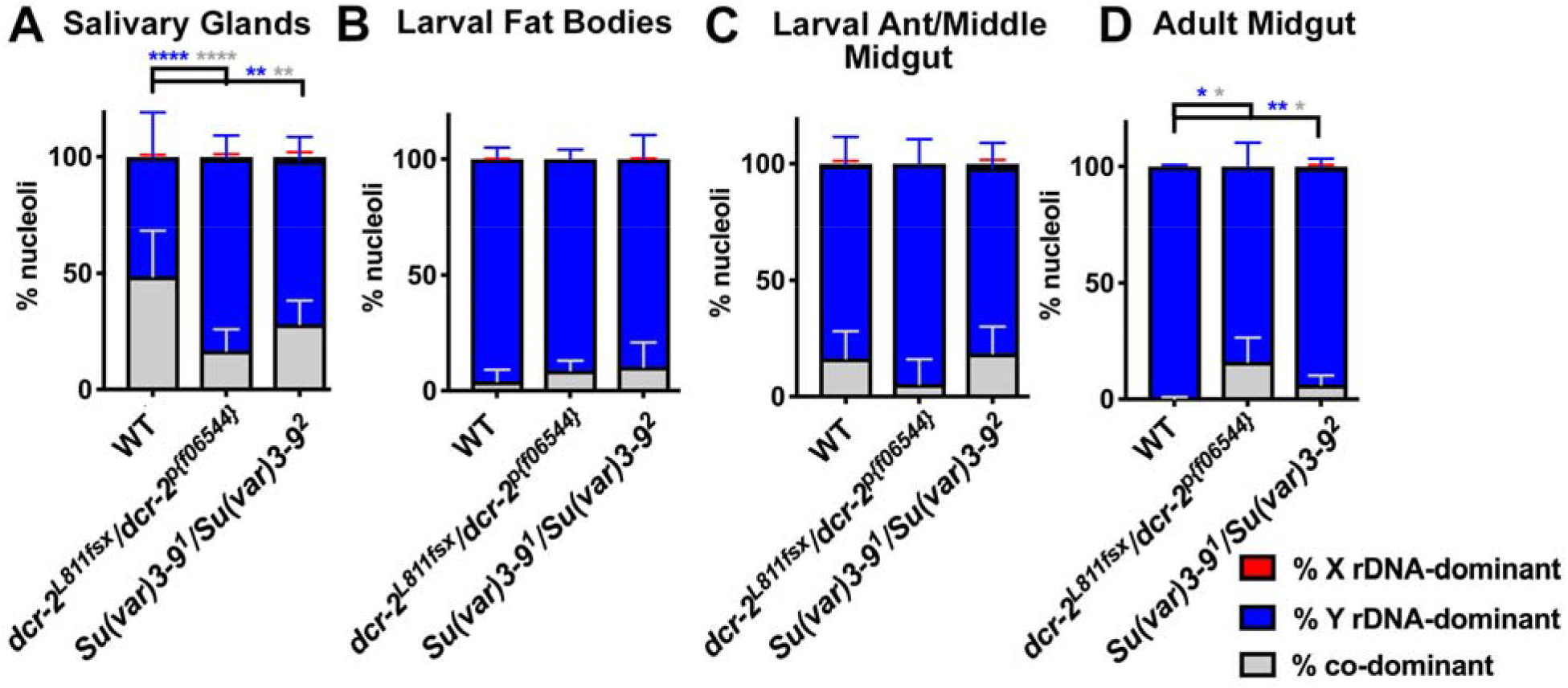
The state of nucleolar dominance in polyploid tissues in *dcr-2* and Su(var)3-9 mutants. Quantification of nucleolar dominance in males in A) salivary glands of wild type (*yw*) (n = 878 cells from 15 salivary glands), *dcr-2^L811fsx^/ dcr-2^p[f06544]^* mutants (n = 373 cells from 8 salivary glands), and *Su(var)3-9^1^/Su(var)3-9^2^* mutants (n = 423 cells from 11 salivary glands), B) larval fat bodies of wild type (n = 1575 cells from 17 fat bodies), *dcr-2^L811fsx^/ dcr-2^p[f06544]^* mutants (n = 320 cells from 4 fat bodies), and *Su(var)3-9^1^/Su(var)3-9^2^* mutants (n = 351 cells from 7 fat bodies), C) larval anterior midgut enterocytes of wild type (n = 181 cells from 6 guts), *dcr-2^L811fsx^/ dcr-2^p[f06544]^* mutants (n = 150 cells from 5 guts), and *Su(var)3-9^1^/Su(var)3-9^2^* mutants (n = 172 cells from 5 guts), and D) adult anterior midgut enterocytes of wild type (n = 922 cells from 7 guts), *dcr-2^L811fsx^/ dcr-2^p[f06544]^* mutants (n = 614 cells from 6 guts), and *Su(var)3-9^1^/Su(var)3-9^2^* mutants (n = 476 cells from 6 guts). Red = % X rDNA-dominant, blue = % Y rDNA-dominant, grey = % co-dominant nuclei. p-values calculated using Welch’s unpaired, unequal variances *t*-test using n = number of tissues. (no star) = not significant, * = < 0.05, ** = < 0.01, **** = < 0.0001. Colors of asterisks correspond to colors of bars for which p-values were calculated (e.g. blue asterisk for Y rDNA-dominant p-values).

**Figure S2:**
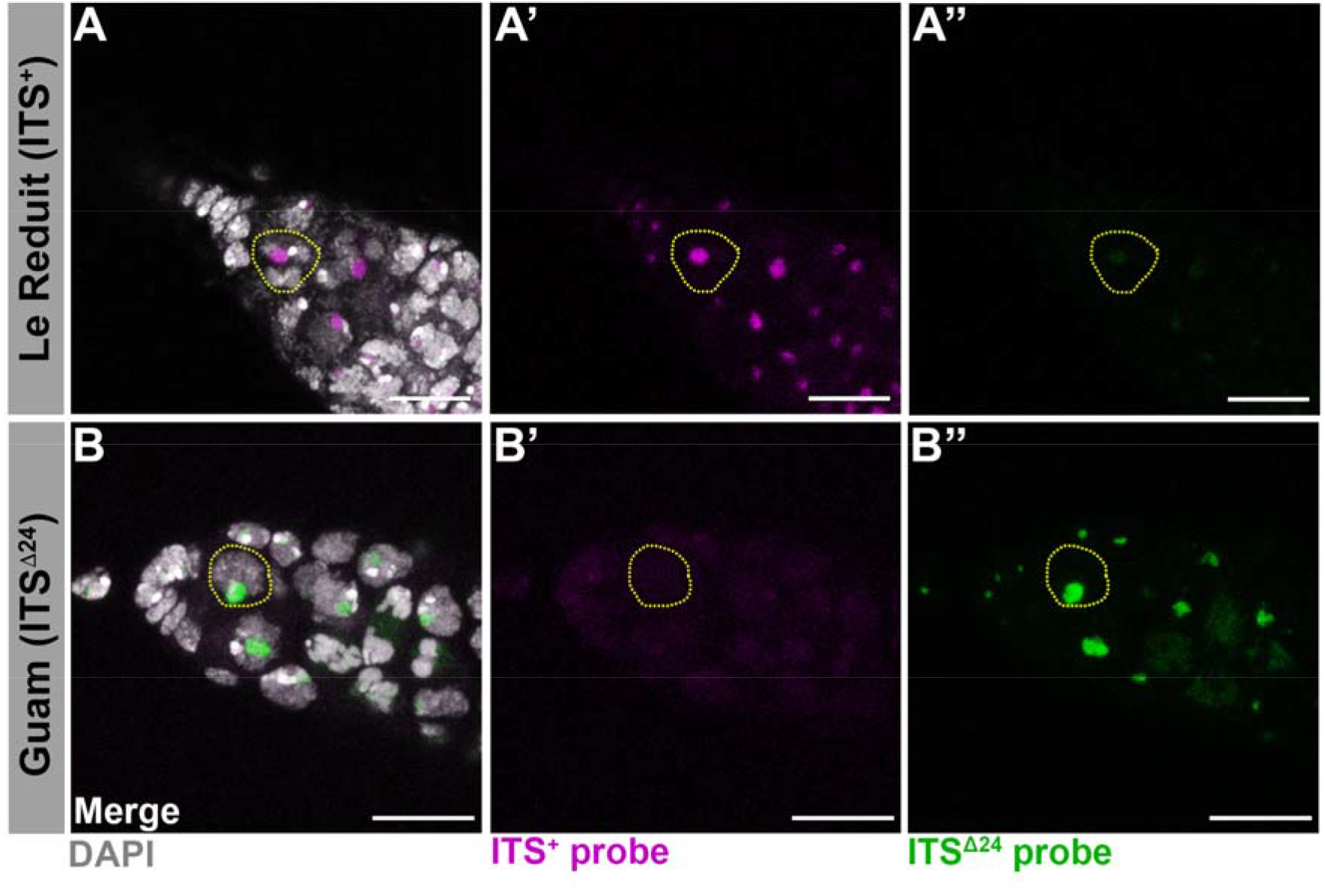
ITS probes that differentially identify distinct X rDNA variants. Representative images of germarium from Le Réduit, and B) Guam *D. melanogaster* strains. All scales = 10μm. Magenta = ITS^+^ rDNA transcript, green = ITS^Δ24^ rDNA transcript, white = DAPI.

**Figure S3:**
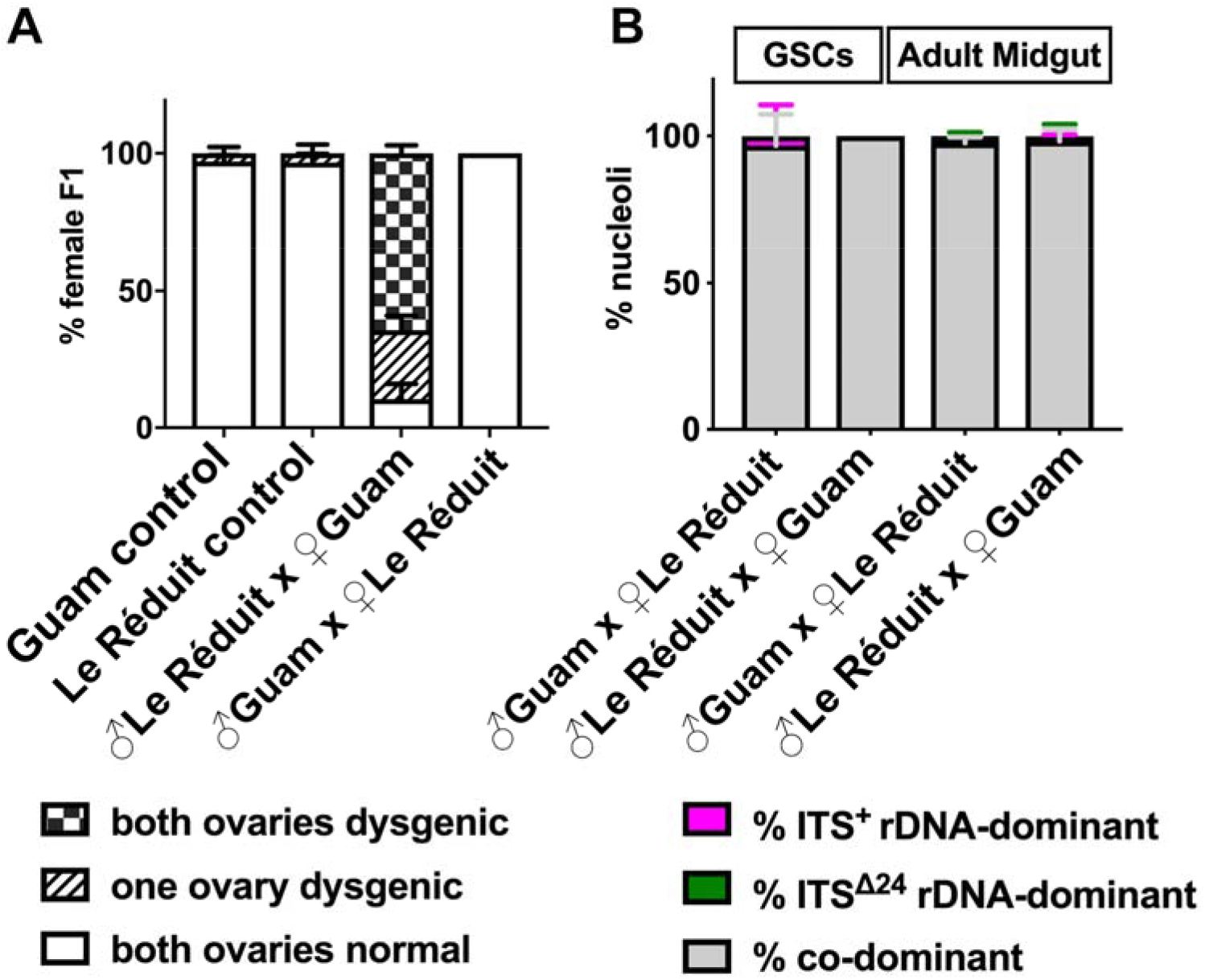
Two X chromosomes (ITS^+^ vs. ITS^Δ24^ variants) exhibit co-dominance irrespective of parental origins. A) Quantification of hybrid dysgenesis in the crosses of 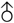 Guam x ♀ Guam controls (n = 104 females from 4 vials), 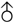 Le Réduit x ♀ Le Réduit controls (n = 185 females from 4 vials), and 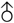 Le Réduit x ♀ Guam (n = 145 females from 3 vials), and 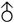 Guam x ♀ Le Réduit (n = 202 females from 4 vials) performed at 25°C (Engels and Preston 1979). B) Quantification of female nucleolar dominance between two X rDNA comparing both cross directions (data from Figure 4 is reproduced to show that the parental origin minimally influence the state of nucleolar dominance). GSCs scored: 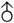 Guam x ♀ Le Réduit (n = 150 cells from 57 germarium), 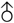 Le Réduit x ♀ Guam (n = 107 cells from 51 germarium). Adult anterior midgut scored: 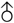 Guam x ♀ Le Réduit (n = 904 cells from 9 guts), 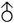 Le Réduit x ♀ Guam (n = 962 cells from 9 guts).

**Figure S4:**
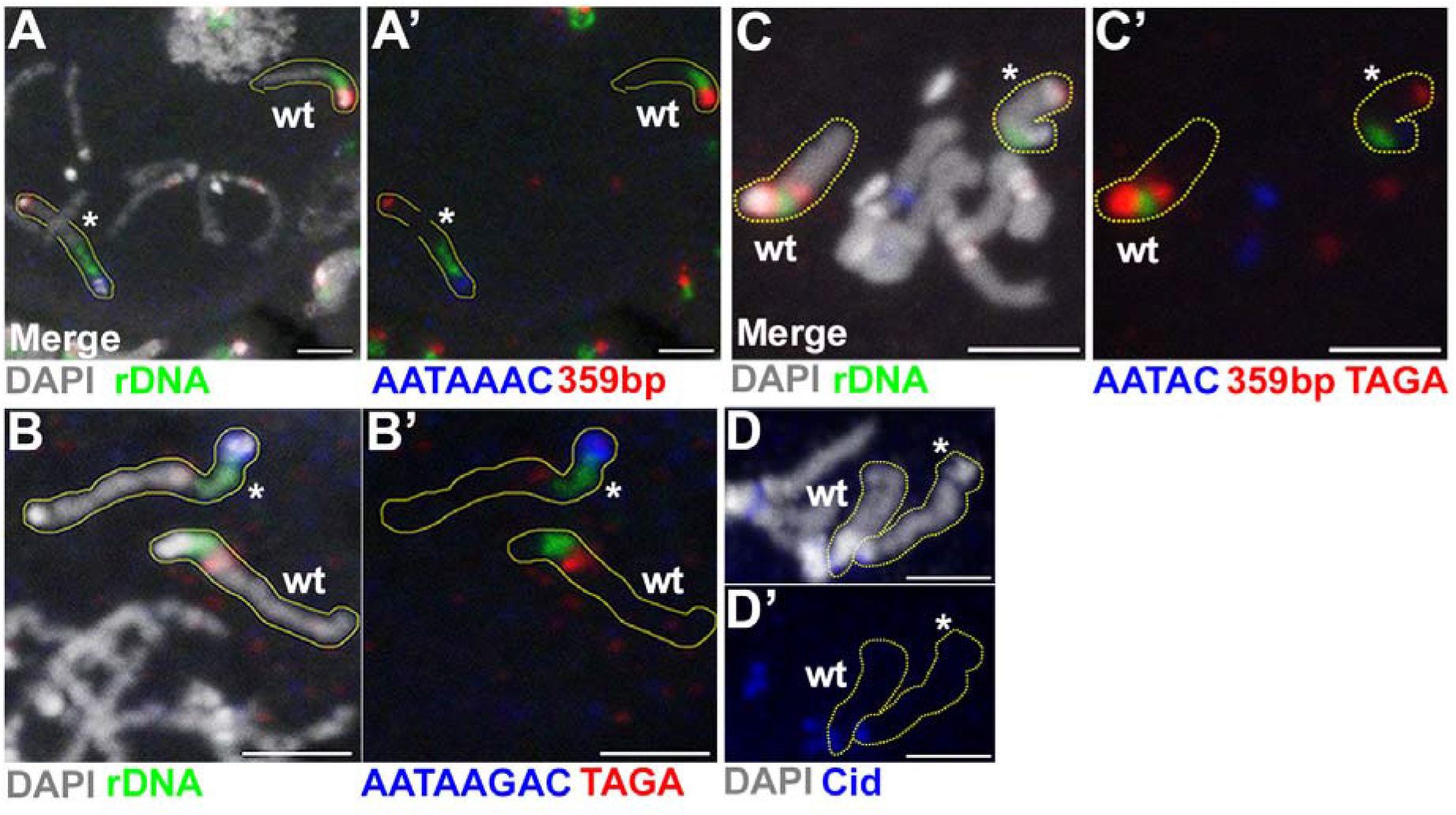
Cytological characterization of the x^bb−^(Y^bb+^) chromosome. DNA fluorescent *in situ* hybridization (FISH) on mitotic chromosome spreads from larval brains of X^bb−^(Y^bb+^)/X females with A) AATAAAC (blue) and 359-bp (red). B) AATAAGAC (blue) and TAGA (red). C) AATAC (blue), TAGA, and 359-bp. D) Immunofluorescence (IF) of centromeric protein Cid (blue) on mitotic chromosome spreads. X^bb−^(Y^bb+^) (*) and wild type X (wt), both outlined in yellow. White = DAPI. All scale bars = 3μm.

## References

Aldrich, J. C. and K. A. Maggert (2015). “Transgenerational inheritance of diet-induced genome rearrangements in Drosophila.” PLoS Genet 11(4): e1005148.

Blum, J. A., S. Bonaccorsi, M. Marzullo, V. Palumbo, Y. M. Yamashita, D. A. Barbash and M. Gatti (2017). “The Hybrid Incompatibility Genes Lhr and Hmr Are Required for Sister Chromatid Detachment During Anaphase but Not for Centromere Function.” Genetics 207(4): 1457–1472.

Bridges, C. B. (1916). “Non-Disjunction as Proof of the Chromosome Theory of Heredity.” Genetics 1(1): 1–52.

Bridges, C. B. (1921). “TRIPLOID INTERSEXES IN DROSOPHILA MELANOGASTER.” Science 54(1394): 252–254.

Busch, H., Y. Daskal, F. Gyorkey and K. Smetana (1979). “Silver staining of nucleolar granules in tumor cells.” Cancer Res 39(3): 857–863.

Cassidy, D. M. and A. W. Blackler (1974). “Repression of nucleolar organizer activity in an interspecific hybrid of the genus Xenopus.” Dev Biol 41(1): 84–96.

Caudy, A. A. and C. S. Pikaard (2002). “Xenopus ribosomal RNA gene intergenic spacer elements conferring transcriptional enhancement and nucleolar dominance-like competition in oocytes.” J Biol Chem 277(35): 31577–31584.

Chandrasekhara, C., G. Mohannath, T. Blevins, F. Pontvianne and C. S. Pikaard (2016). “Chromosome-specific NOR inactivation explains selective rRNA gene silencing and dosage control in Arabidopsis.” Genes Dev 30(2): 177–190.

Chen, Z. J., L. Comai and C. S. Pikaard (1998). “Gene dosage and stochastic effects determine the severity and direction of uniparental ribosomal RNA gene silencing (nucleolar dominance) in Arabidopsis allopolyploids.” Proc Natl Acad Sci U S A 95(25): 14891–14896.

Chen, Z. J. and C. S. Pikaard (1997). “Transcriptional analysis of nucleolar dominance in polyploid plants: biased expression/silencing of progenitor rRNA genes is developmentally regulated in Brassica.” Proc Natl Acad Sci U S A 94(7): 3442–3447.

Coen, E. S. and G. A. Dover (1982). “Multiple Pol I initiation sequences in rDNA spacers of Drosophila melanogaster.” Nucleic Acids Res 10(21): 7017–7026.

Costa-Nunes, P., O. Pontes, S. B. Preuss and C. S. Pikaard (2010). “Extra views on RNA-dependent DNA methylation and MBD6-dependent heterochromatin formation in nucleolar dominance.” Nucleus 1(3): 254–259.

Croce, C. M., A. Talavera, C. Basilico and O. J. Miller (1977). “Suppression of production of mouse 28S ribosomal RNA in mouse-human hybrids segregating mouse chromosomes.” Proc Natl Acad Sci U S A 74(2): 694–697.

Durica, D. S. and H. M. Krider (1977). “Studies on the ribosomal RNA cistrons in interspecific Drosophila hybrids. I. Nucleolar dominance.” Dev Biol 59(1): 62–74.

Earley, K., R. J. Lawrence, O. Pontes, R. Reuther, A. J. Enciso, M. Silva, N. Neves, M. Gross, W. Viegas and C. S. Pikaard (2006). “Erasure of histone acetylation by Arabidopsis HDA6 mediates large-scale gene silencing in nucleolar dominance.” Genes Dev 20(10): 1283–1293.

Earley, K. W., F. Pontvianne, A. T. Wierzbicki, T. Blevins, S. Tucker, P. Costa-Nunes, O. Pontes and C. S. Pikaard (2010). “Mechanisms of HDA6-mediated rRNA gene silencing: suppression of intergenic Pol II transcription and differential effects on maintenance versus siRNA-directed cytosine methylation.” Genes Dev 24(11): 1119–1132.

Engels, W. R. and C. R. Preston (1979). “Hybrid dysgenesis in Drosophila melanogaster: the biology of female and male sterility.” Genetics 92(1): 161–174.

Felle, M., J. H. Exler, R. Merkl, K. Dachauer, A. Brehm, I. Grummt and G. Langst (2010). “DNA sequence encoded repression of rRNA gene transcription in chromatin.” Nucleic Acids Res 38(16): 5304–5314.

Ghoshal, K., S. Majumder, J. Datta, T. Motiwala, S. Bai, S. M. Sharma, W. Frankel and S. T. Jacob (2004). “Role of human ribosomal RNA (rRNA) promoter methylation and of methyl-CpG-binding protein MBD2 in the suppression of rRNA gene expression.” J Biol Chem 279(8): 6783–6793.

Greil, F. and K. Ahmad (2012). “Nucleolar dominance of the Y chromosome in Drosophila melanogaster.” Genetics 191(4): 1119–1128.

Grewal, S. S., L. Li, A. Orian, R. N. Eisenman and B. A. Edgar (2005). “Myc-dependent regulation of ribosomal RNA synthesis during Drosophila development.” Nat Cell Biol 7(3): 295–302.

Grimaldi, G., P. Fiorentini and P. P. Di Nocera (1990). “Spacer promoters are orientation-dependent activators of pre-rRNA transcription in Drosophila melanogaster.” Mol Cell Biol 10(9): 4667–4677.

Heix, J. and I. Grummt (1995). “Species specificity of transcription by RNA polymerase I.” Curr Opin Genet Dev 5(5): 652–656.

Heliot, L., F. Mongelard, C. Klein, M. F. O’Donohue, J. M. Chassery, M. Robert-Nicoud and Y. Usson (2000). “Nonrandom distribution of metaphase AgNOR staining patterns on human acrocentric chromosomes.” J Histochem Cytochem 48(1): 13–20.

Houchins, K., M. O’Dell, R. B. Flavell and J. P. Gustafson (1997). “Cytosine methylation and nucleolar dominance in cereal hybrids.” Mol Gen Genet 255(3): 294–301.

Jagannathan, M., N. Warsinger-Pepe, G. J. Watase and Y. M. Yamashita (2017). “Comparative Analysis of Satellite DNA in the Drosophila melanogaster Species Complex.” G3 (Bethesda) 7(2): 693–704.

Kidd, S. J. and D. M. Glover (1981). “Drosophila melanogaster ribosomal DNA containing type II insertions is variably transcribed in different strains and tissues.” J Mol Biol 151(4): 645–662.

Labhart, P. and R. H. Reeder (1984). “Enhancer-like properties of the 60/81 bp elements in the ribosomal gene spacer of Xenopus laevis.” Cell 37(1): 285–289.

Larracuente, A. M. and P. M. Ferree (2015). “Simple method for fluorescence DNA in situ hybridization to squashed chromosomes.” J Vis Exp(95): 52288.

Lawrence, R. J., K. Earley, O. Pontes, M. Silva, Z. J. Chen, N. Neves, W. Viegas and C. S. Pikaard (2004). “A concerted DNA methylation/histone methylation switch regulates rRNA gene dosage control and nucleolar dominance.” Mol Cell 13(4): 599–609.

Levesque, M. J., P. Ginart, Y. Wei and A. Raj (2013). “Visualizing SNVs to quantify allele-specific expression in single cells.” Nat Methods 10(9): 865–867.

Long, E. O. and I. B. Dawid (1980). “Repeated genes in eukaryotes.” Annu Rev Biochem 49: 727–764.

Lu, K. L., J. O. Nelson, G. J. Watase, N. Warsinger-Pepe and Y. M. Yamashita (2018). “Transgenerational dynamics of rDNA copy number in Drosophila male germline stem cells.” Elife 7.

Macleod, D. and A. Bird (1982). “DNAase I sensitivity and methylation of active versus inactive rRNA genes in xenopus species hybrids.” Cell 29(1): 211–218.

Mayer, C., K. M. Schmitz, J. Li, I. Grummt and R. Santoro (2006). “Intergenic transcripts regulate the epigenetic state of rRNA genes.” Mol Cell 22(3): 351–361.

Mohannath, G., F. Pontvianne and C. S. Pikaard (2016). “Selective nucleolus organizer inactivation in Arabidopsis is a chromosome position-effect phenomenon.” Proc Natl Acad Sci U S A 113(47): 13426–13431.

Moss, T. and V. Y. Stefanovsky (2002). “At the center of eukaryotic life.” Cell 109(5): 545–548.

Murayama, A., K. Ohmori, A. Fujimura, H. Minami, K. Yasuzawa-Tanaka, T. Kuroda, S. Oie, H. Daitoku, M. Okuwaki, K. Nagata, A. Fukamizu, K. Kimura, T. Shimizu and J. Yanagisawa (2008). “Epigenetic control of rDNA loci in response to intracellular energy status.” Cell 133(4): 627–639.

Neves, N., J. S. Heslop-Harrison and W. Viegas (1995). “rRNA gene activity and control of expression mediated by methylation and imprinting during embryo development in wheat x rye hybrids.” Theor Appl Genet 91(3): 529–533.

Peng, J. C. and G. H. Karpen (2007). “H3K9 methylation and RNA interference regulate nucleolar organization and repeated DNA stability.” Nat Cell Biol 9(1): 25–35.

Pikaard, C. S. (2000). “Nucleolar dominance: uniparental gene silencing on a multi-megabase scale in genetic hybrids.” Plant Mol Biol 43(2–3): 163–177.

Pontes, O., R. J. Lawrence, M. Silva, S. Preuss, P. Costa-Nunes, K. Earley, N. Neves, W. Viegas and C. S. Pikaard (2007). “Postembryonic establishment of megabase-scale gene silencing in nucleolar dominance.” PLoS One 2(11): e1157.

Pontes, O., C. F. Li, P. Costa Nunes, J. Haag, T. Ream, A. Vitins, S. E. Jacobsen and C. S. Pikaard (2006). “The Arabidopsis chromatin-modifying nuclear siRNA pathway involves a nucleolar RNA processing center.” Cell 126(1): 79–92.

Pontes, O., N. Neves, M. Silva, M. S. Lewis, A. Madlung, L. Comai, W. Viegas and C. S. Pikaard (2004). “Chromosomal locus rearrangements are a rapid response to formation of the allotetraploid Arabidopsis suecica genome.” Proc Natl Acad Sci U S A 101(52): 18240–18245.

Pontvianne, F., T. Blevins, C. Chandrasekhara, W. Feng, H. Stroud, S. E. Jacobsen, S. D. Michaels and C. S. Pikaard (2012). “Histone methyltransferases regulating rRNA gene dose and dosage control in Arabidopsis.” Genes Dev 26(9): 945–957.

Preuss, S. and C. S. Pikaard (2007). “rRNA gene silencing and nucleolar dominance: insights into a chromosome-scale epigenetic on/off switch.” Biochim Biophys Acta 1769(5–6): 383–392.

Preuss, S. B., P. Costa-Nunes, S. Tucker, O. Pontes, R. J. Lawrence, R. Mosher, K. D. Kasschau, J. C. Carrington, D. C. Baulcombe, W. Viegas and C. S. Pikaard (2008). “Multimegabase silencing in nucleolar dominance involves siRNA-directed DNA methylation and specific methylcytosine-binding proteins.” Mol Cell 32(5): 673–684.

Probst, A. V., M. Fagard, F. Proux, P. Mourrain, S. Boutet, K. Earley, R. J. Lawrence, C. S. Pikaard, J. Murfett, I. Furner, H. Vaucheret and O. Mittelsten Scheid (2004). “Arabidopsis histone deacetylase HDA6 is required for maintenance of transcriptional gene silencing and determines nuclear organization of rDNA repeats.” Plant Cell 16(4): 1021–1034.

Reeder, R. H., J. G. Roan and M. Dunaway (1983). “Spacer regulation of Xenopus ribosomal gene transcription: competition in oocytes.” Cell 35(2 Pt 1): 449–456.

Roussel, P., C. Andre, L. Comai and D. Hernandez-Verdun (1996). “The rDNA transcription machinery is assembled during mitosis in active NORs and absent in inactive NORs.” J Cell Biol 133(2): 235–246.

Santoro, R. and I. Grummt (2005). “Epigenetic mechanism of rRNA gene silencing: temporal order of NoRC-mediated histone modification, chromatin remodeling, and DNA methylation.” Mol Cell Biol 25(7): 2539–2546.

Schlotterer, C., M. T. Hauser, A. von Haeseler and D. Tautz (1994). “Comparative evolutionary analysis of rDNA ITS regions in Drosophila.” Mol Biol Evol 11(3): 513–522.

Schmitz, K. M., C. Mayer, A. Postepska and I. Grummt (2010). “Interaction of noncoding RNA with the rDNA promoter mediates recruitment of DNMT3b and silencing of rRNA genes.” Genes Dev 24(20): 2264–2269.

Smetana, K. and H. Busch (1964). “STUDIES ON THE ULTRASTRUCTURE OF THE NUCLEOLI OF THE WALKER TUMOR AND RAT LIVER.” Cancer Res 24: 537–557.

Tautz, D., J. M. Hancock, D. A. Webb, C. Tautz and G. A. Dover (1988). “Complete sequences of the rRNA genes of Drosophila melanogaster.” Mol Biol Evol 5(4): 366–376.

Tautz, D., C. Tautz, D. Webb and G. A. Dover (1987). “Evolutionary divergence of promoters and spacers in the rDNA family of four Drosophila species. Implications for molecular coevolution in multigene families.” J Mol Biol 195(3): 525–542.

Tucker, S., A. Vitins and C. S. Pikaard (2010). “Nucleolar dominance and ribosomal RNA gene silencing.” Curr Opin Cell Biol 22(3): 351–356.

Wilk, R., S. U. M. Murthy, H. Yan and H. M. Krause (2010). “In Situ Hybridization: Fruit Fly Embryos and Tissues.” Current Protocols Essential Laboratory Techniques 4(1): 9.3.1–9.3.24.

Zhou, J., T. B. Sackton, L. Martinsen, B. Lemos, T. H. Eickbush and D. L. Hartl (2012). “Y chromosome mediates ribosomal DNA silencing and modulates the chromatin state in Drosophila.” Proc Natl Acad Sci U S A 109(25): 9941–9946.

